# Longitudinal analysis of the B-cell receptor repertoire across 30 years of ageing

**DOI:** 10.64898/2026.08.18.745470

**Authors:** Lois Leenders, Maud A.J.I. van den Oetelaar, Peter Engelfriet, Anne-Marie Buisman, Mary-Lène de Zeeuw-Brouwer, Lia de Rond, W.M. Monique Verschuren, Roel C.H. Vermeulen, Anton W. Langerak, P. Martijn Kolijn

## Abstract

**Background:** The gradual decline in the functionality of the immune system during aging is commonly referred to as immunosenescence. This study aims to investigate changes in the B-cell receptor immunoglobulin heavy chain (BCR IGH) gene repertoire during natural aging and evaluate the dynamics of emergent low-level BCR IGH clonality in the elderly. We conduct a longitudinal study nested within the Doetinchem Cohort study, comprising 98 participants aged between 31 and 59 years old at study entry who had repeated blood samples drawn at 5 year intervals over a 30 year period (n=548 samples). We sequenced the IGH gene repertoire and evaluated the impact of aging on IGH gene repertoire clonality and diversity using linear mixed effects modeling.

**Results:** Participants older than 60 years exhibited increased BCR IGH clonality and reduced IGH gene repertoire diversity. In a multivariable model, IGH gene repertoire diversity was significantly decreased for individuals with a dominant clonotype ratio greater than 10 (β =-0.57, p < 0.001). Additionally, a trend toward reduced IGH gene repertoire diversity was observed in participants aged 60-70 years (β =-0.19, p = 0.1) and those aged 70 years or older ( β =-0.20, p = 0.13). IGH gene repertoire diversity was determined primarily by the naïve and transitional B-cell pool, while BCR IGH clonality was influenced by switched memory and age-associated B-cell counts.

**Conclusions:** In summary, our study indicates that IGH gene repertoire diversity decreases significantly after age 60, which coincides with an increased incidence of low-level BCR IGH clonality. This clonality may be driven largely by switched memory and age-associated B-cells. By providing deep insights into age-related dynamic changes in the IGH gene repertoire, these findings lay the groundwork for the molecular assessment and monitoring of incident clonality by clinicians and researchers alike.

## Introduction

Increasing human life expectancy has led to a rapidly expanding aging population and a corresponding rise in age-related diseases, including cancer and severe or chronic infections.^1^ This susceptibility is caused by a gradual decline in the functionality of the immune system, commonly referred to as immunosenescence.^2^ Immunosenescence encompasses both structural and functional alterations within the immune system, including remodeling of immune organs and dysregulation of innate and adaptive immune responses. A hallmark of this process is chronic low-grade inflammation, resulting from inadequate antigen clearance, persistence of senescent cells with a ‘senescence-associated phenotype’ (SASP), increased levels of pro-inflammatory cytokines, chemokines and reactive oxygen species, and selected stressors (*e*.*g*. UV and gamma radiation, environmental exposures).^3^ Another prominent feature is thymic involution, the age-related shrinking and functional decline of the thymus.^2^ As a result of these factors, the T-cells of elderly individuals are replenished at a lower rate and tend to be hyporesponsive to antigen stimulation.^2^

In parallel with these T-cell alterations, B-cell immunity also undergoes substantial age-related changes.^4^ Ageing is associated with reduced antibody production, and decreased antibody affinity. In addition, defects in B cell tolerance mechanisms are more common in elderly individuals, contributing to the production of autoreactive antibodies.^5^ Age-related changes also affect B cell development, which reduces B cell production in the bone marrow and the overall number of mature human B cells.^4^ These systemic changes disrupt B cell homeostasis, resulting in an altered composition and functionality of the B cell compartment over time.^6^

A central determinant of B-cell function is the B-cell receptor (BCR) repertoire, which enables recognition of a wide variety of antigens. The BCR is a membrane-bound immunoglobulin (IG) comprising two identical heavy chains and two identical light chains.^7^ The variable region of the immunoglobulin heavy chain (IGH) is encoded by randomly combined variable (V), diversity (D), and joining (J) genes, while the light chain is encoded by randomly combined V and J genes. Together, the heavy chain and light chain combine to create a BCR unique to a cell or to a clone of cells.^8^ The collection of all these unique receptors constitutes the BCR repertoire. BCR IGH gene repertoire diversity is essential to enable an effective immune response to the broad range of pathogens that individuals will encounter during their lifespan.

A prominent feature of immune aging is the emergence of clonal B-cell expansions, reflected by reduced BCR repertoire diversity.^9^ One well-recognized manifestation of this phenomenon is monoclonal B-cell lymphocytosis (MBL), defined by the presence of clonal B cells in the peripheral blood of otherwise healthy individuals.^10^ MBL is a common condition among individuals above 50 years old (5-10% of the healthy population) and exceedingly common among the elderly (up to 50% of individuals over 90 years old).^11^ Although MBL is an indolent condition, individuals with MBL are at increased risk of progression to chronic lymphocytic leukemia (CLL), serious infections, and non-hematological malignancies as well as have reduced humoral immune response to vaccinations.^12^

Despite growing recognition of age-related changes in B-cell clonality and repertoire diversity, most studies examining IGH gene repertoire alterations during aging have relied on cross-sectional designs. Consequently, the long-term dynamics of repertoire diversity and clonal expansions within individuals during natural aging remain poorly understood.

In this study, we investigated age-related changes in the IGH gene repertoire using a unique longitudinal dataset spanning 30 years, consisting of buffy coat samples collected at five-year intervals from 98 healthy adults participating in the Doetinchem Cohort Study. By analyzing IGH gene repertoire diversity and clonal dynamics over time, we aimed to characterize how IGH gene repertoire diversity evolves during aging and to evaluate the emergence and persistence of low-level B-cell clonality in the general population.

## Methods

### Study design

Our study was nested within the Doetinchem Cohort study, a population study centered around the Dutch town Doetinchem **(Table 1)**.^13,14^ The Doetinchem Cohort Study was designed to study the longitudinal impact of lifestyle factors and biological risk factors on the incidence of chronic diseases, physical and cognitive functioning and quality of life. We included equal numbers of men and women. Median age at study entry was 39 years (range 31-69 years). We included 548 repeated samples from 98 individuals drawn at 5 year intervals over a 30 year period. The median number of repeated samples available per individual was 6 (range 3-7). Flow cytometry data were available at sampling round 6 and 7 only. Viral titers for Epstein-Barr virus (EBV), Cytomegalovirus (CMV) and Varicella Zoster virus (VZV) were available at round 6 only.

**Table 1.**
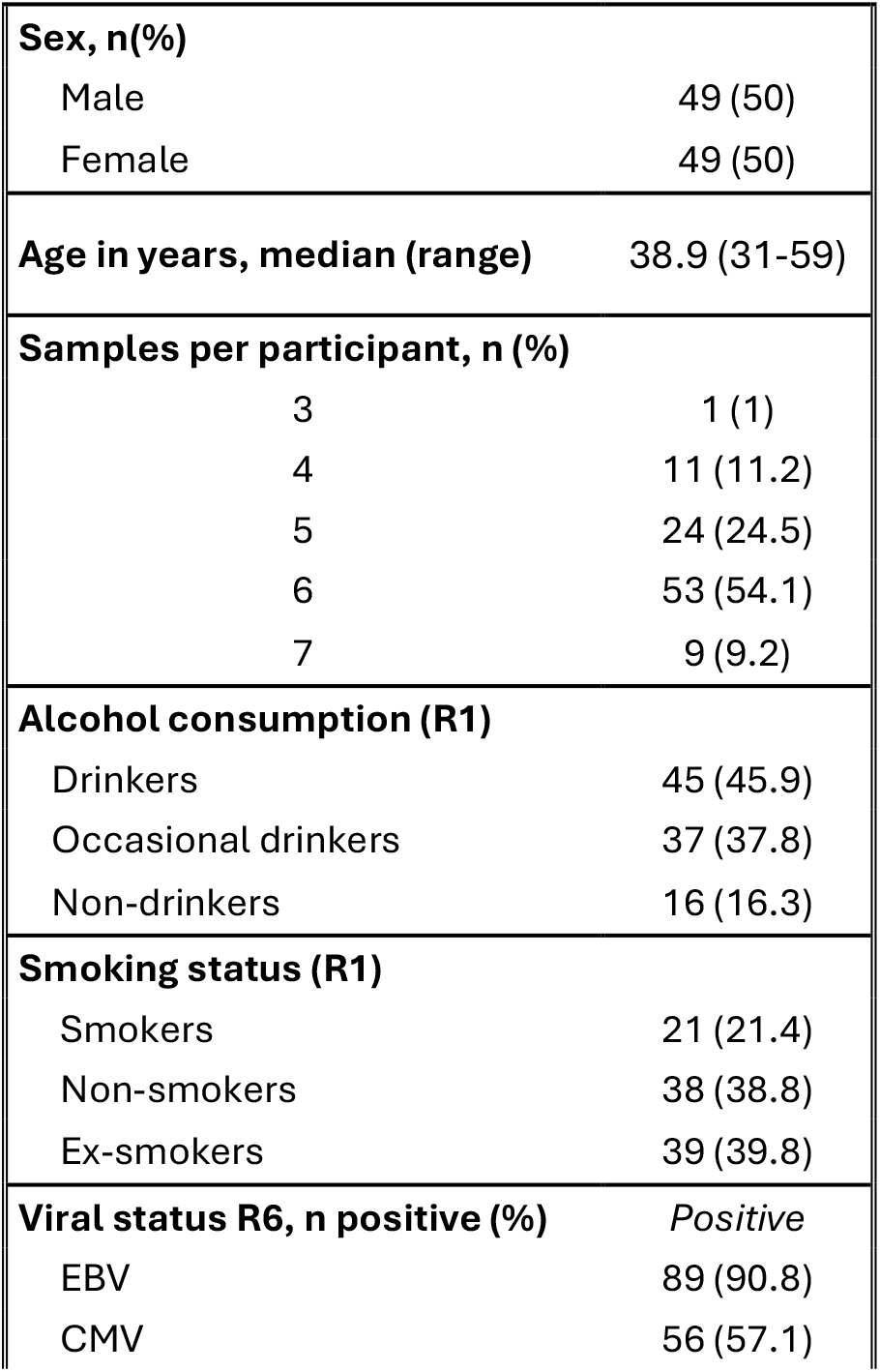
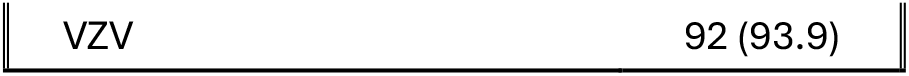
Baseline characteristics of the study population. Data on alcohol consumption and smoking status data are from sampling round 1 (R1), and data on seropositivity for Epstein-Barr virus (EBV), Cytomegalovirus (CMV), and Varicella Zoster virus (VZV) are from sampling round 6 (R6). Age is given at study entry, meaning R1. Occasional drinkers: mean intake <1 glass of alcohol per day. Drinkers: mean intake > 1 glass of alcohol per day.

### Next-generation sequencing

We used 500 ng of buffy coat DNA as input for the leader-based multiplex PCR of IGHV-IGHD-IGHJ rearrangements.^15^ The following PCR program was run for all samples: denaturation at 95 °C for 3 min; 35 cycles of 95 °C for 45 s, 63 °C for 45 s, 68 °C for 1 min; final extension at 68 °C for 10 min; 12 °C on hold. After amplification, PCR products were purified on Agencourt AMPURE beads (Beckman Coulter, Brea, CA, USA) and eluted using water. The purified product was checked for primer dimers using the D1000 DNA ScreenTape assay on a 4200 Agilent Tapestation (Agilent, Santa Clara, CA, USA). Paired-end sequencing was performed using the MiSeq Reagent Kit v3 (2 × 300 bp) on the MiSeq Benchtop Sequencer (Illumina, San Diego, CA, USA). PhiX was spiked-in at a 10% concentration to increase library diversity. The BCR IGH repertoire was characterized using the ARResT/Interrogate immunoprofiler, an R/Shiny based tool for in silico immunoprofiling developed by the EuroClonality-NGS working group.^16^ ARResT/Interrogate annotates the variable (V), diversity (D) and joining (J) genes for each rearrangement and determines the complementarity-determining region 3 (CDR3) by processing multiple IMGT/HighV-QUEST runs. Clonotypes were computed as unique pairs of IGHV genes and CDR3 amino acid sequences within a given sample. Only reads annotated as productive IGH rearrangements were included. Clonotypes belonging to the same clonal lineage were collapsed. Clonotypes belonging to the same clonal lineage were defined as expressing the same IGHV gene with up to 2 mismatching amino acids in the CDR3. ARResT/Interrogate uses IMGT’s definition of unproductive rearrangements as an out-of-frame junction, due to one or more of the following causes: stop codon(s), frameshift mutation(s), defects in the splicing sites and/or the regulatory element(s), unusual features such as translocation or gene fusion, and/or changes of conserved amino acids demonstrated to lead to incorrect folding.

### Quality control

In every PCR, a central in-tube quality control (cIT-QC) was spiked.^17^ This cIT-QC consists of DNA from 9 selected cell lines bearing 46 rearrangements, 7 of which are IGH rearrangements. A total of 280 IGH copies (2 μL) are spiked in every PCR replicate before amplification, allowing calculation of a conversion factor from reads to cells. This conversion factor can be used to partly correct for differential amplification bias and overamplification in B-cell depleted samples. The cIT-QC was developed by the EuroClonality-NGS working group.^15^ The median number of reads with IGH junction per sample was 223,746 (range 22,004-585,153). After spike-in conversion, this yielded a median template input of 3447 B-cells per sample (range 57-17,361).

### Clonality and IGH gene repertoire diversity evaluation

IGH gene repertoire diversity was evaluated using the Shannon Diversity Index calculated through the diversity function in the “vegan” (2.7.1) R package.^18^ IGH clonality was evaluated using the dominant clonotype ratio.^19^ Dominant clonotype ratio is calculated by dividing the frequency of the most dominant IGH clonotype by the mean of the frequency of the clonotypes ranked 3^rd^-7^th^. Clonotypes with a dominant clonotype ratio between 3 and 10 were designated as “minor clonotypes”, while clonotypes with a dominant clonotype ratio exceeding 10 were designated as “major clonotypes”. The dominant clonotype ratio was log_2_-transformed to preserve low-level clonality dynamics in the figures. Persistent clonality was defined as a minor or major clonotype that remained clonal (i.e. reached the threshold of dominant clonotype ratio >3 for a minor clonotype or >10 for a major clonotype) in at least two sequentially collected samples. Only clonotypes belonging to the same clonal lineage were scored as persistent.

### Flow cytometry

As previously described, fresh whole blood samples from the DCS subcohort participants were collected between August 2016 and March 2017 and were processed and analyzed within 6 hours on a 4-laser LSRII Fortessa X20 flow cytometer (BD Biosciences) for absolute numbers of leukocyte subsets (cell counts µL-1).^20,21^ Two labeled antibody panels per participant were used with a lyse-no-wash protocol, one in a TruCOUNT® tube (BD Biosciences) and one in a common Falcon tube. A previously published gating strategy was used.^20,21^

Repeated blood samples were collected in December 2024 for 62 participants. B-cell counts were again determined using a TruCOUNT® tube (BD Biosciences). Antibody mixes were prepared in FACS buffer consisting of PBS (without Ca/Mg; Gibco as part of Thermo Fisher Scientific, Bleiswijk, NL) with 0.5 % bovine serum albumin (BSA; Sigma-Aldrich, Zwijndrecht, NL) and 2 mM ethylenediaminetetraacetic acid (EDTA; Thermo Fisher Scientific). Additionally, 20 µl Brilliant stain buffer (BD Biosciences, Franklin Lakes, NJ, USA) was added per donor. Next, 50 µl EDTA blood was added, and after mixing, samples were incubated for 30 min at RT in the dark. Afterwards, red blood cells were lysed with a 10x diluted BD FACS lysis buffer (BD Biosciences) that was incubated for 15 min at RT in the dark. Samples were processed and analyzed within 6 hours on a 5-laser Symphony A3 flow cytometer (BD Biosciences) for absolute numbers of leukocyte subsets (cell counts µL-1) according to manufacturer’s instructions. The antibody panel used for the repeated blood samples from 2024 is described in Supplementary Table 1. An important change to the 2017 panel is the addition of CD21 and CD11c to the 2024 panel, enabling differentiation between double negative (DN, CD27^-^IgD^-^) B-cell subsets. The most relevant DN B-cell subset is DN2 (CD21^-^CD11c^+^), as this subset represents a group of B-cells associated with aging and autoimmunity.^22^

### Serology

As previously described, Anti-CMV and anti-EBV antibody levels were measured in plasma samples of the study participants with an in-house developed multiplex immuno-assay (MIA).^21,23^ Individuals were CMV-seropositive with a level of more than 5 relative units (RU)/ml, whereas the threshold for EBV seropositivity was 22 RU/ml and individuals were considered EBV-seronegative with a level ≤ 16 RU/ml.^23^

### Statistical analysis

All statistical analyses and data plotting were conducted using R software version 4.5.1 (R Core Team, 2025).^24^ Scatter and spaghetti plots were created with the ggplot2 package.^25^ Visualization of the relationship between diversity and age was performed using Locally Estimated Scatterplot Smoothing (LOESS) regression using the “geom_smooth” function in the ggplot2 R package. Linear mixed effects model 1 included IGH gene repertoire diversity as the outcome variable and 10-year age category (30-40, 40-50, 50-60, 60-70 and 70+), sex, smoking status and alcohol consumption as fixed effects and subject ID as a random effect. Linear mixed effects model 2 included log2-transformed dominant clonotype ratio as the outcome variable and 10-year age category (30-40, 40-50, 50-60, 60-70 and 70+), sex, smoking status and alcohol consumption as fixed effects and subject ID as a random effect. In model 3 for IGH gene repertoire diversity, we added clonality (no clonality, minor clonotype, major clonotype) as a fixed effect to evaluate whether the decreased diversity in the higher age categories was driven by monoclonality or a broader reduction of repertoire diversity.

Model 4 included B-cell count per microliter blood as the dependent variable, IGH gene repertoire diversity, log2-transformed dominant clonotype ratio, age in years, sex and smoking status and alcohol consumption as fixed effects and subject ID as a random effect. B-cell count data was only available for round 6 and 7 of the study, meaning the B-cell count analyses are limited to these two rounds only. Mixed effects models were fitted using the “lmer” function from the “lme4” (1.1.37) R package.^26^ For the mixed effects models, p-values were calculated using the “lmerTest” package, which adjusts the degrees of freedom using Satterthwaite’s (Kenward-Roger’s) approximations.^27^ Threshold for significance was placed at a p-value of 0.05. As viral titer measurements were only available for round 6 of the study, we employed a logistic regression model (model 5) to evaluate the relationship between clonality category (dominant clonotype ratio above or below 10) and CMV, EBV and VZV titer, adjusted for age and sex. Model 5 was fitted using the “glm” function from the “stats” (3.6.2) R package.

## Results

### Effect of age on IGH gene repertoire diversity

Using Locally Estimated Scatterplot Smoothing (LOESS) regression, we observed a decrease in IGH gene repertoire diversity starting around age 60 (Figure 1). We employed a linear mixed-effects model adjusted for sex, smoking status, and alcohol consumption to quantify the effect of 10-year age categories on IGH gene repertoire diversity. The model revealed that IGH gene repertoire diversity was significantly decreased for participants in age categories 60-70 (β =-0.24, p = 0.02) and 70+ (β =-0.27, p < 0.0001, Supplementary Table 2).

**Figure 1.**
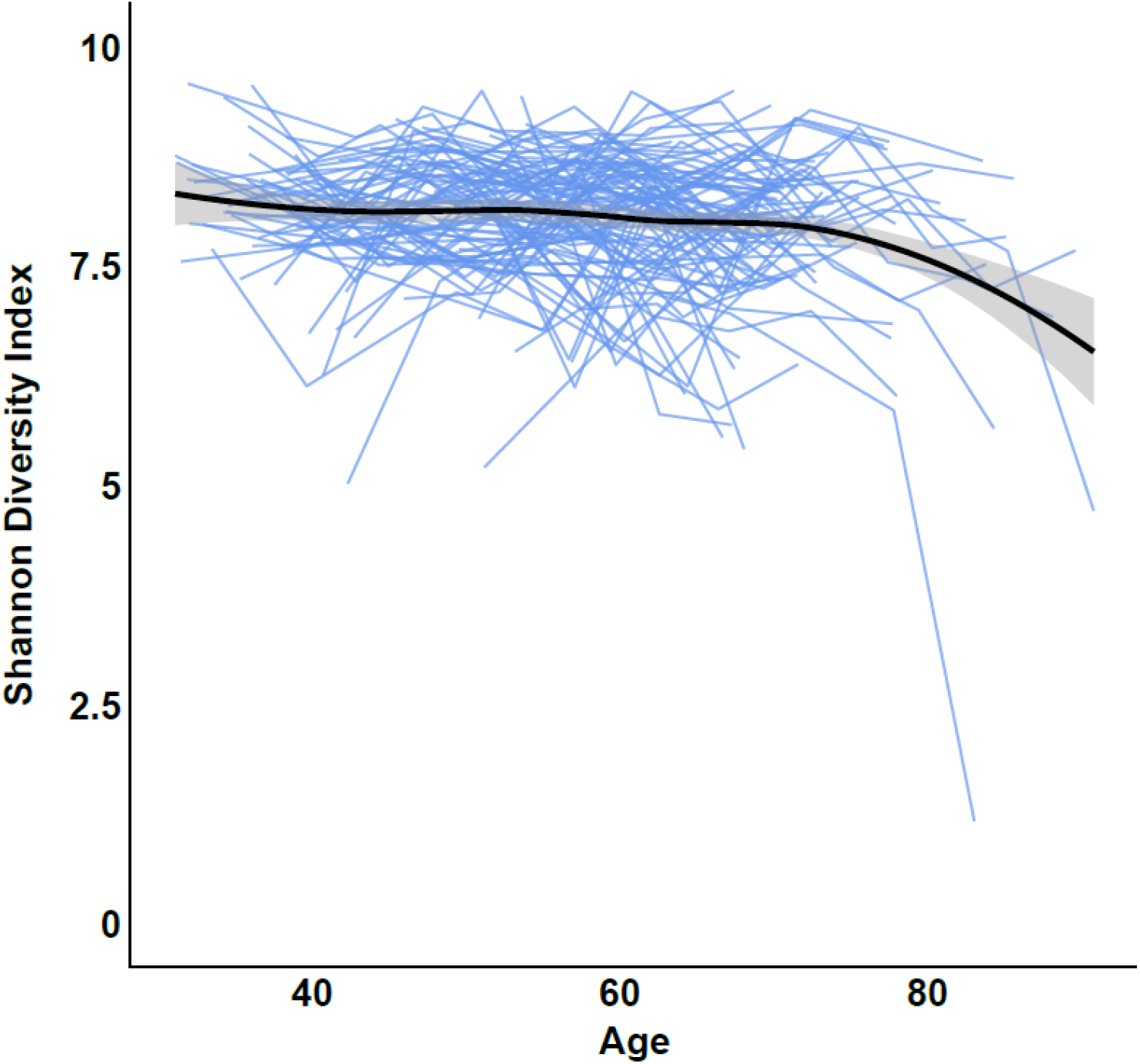
The relationship between age and IGH gene repertoire diversity. Spaghetti plot with blue lines representing repeated measurements from single individuals. The black line represents the Locally Estimated Scatterplot Smoothing (LOESS) regression estimate that summarizes the general trend. X-axis: age in years, Y-axis: Shannon Diversity Index score for IGH gene repertoire diversity. Number of observations: 548. Number of individuals: 98.

### Evolution of B-cell clonality patterns in healthy adults

We evaluated BCR IGH clonality in 548 repeated buffy coat samples from the 98 adults in our study (Figure 2). Forty-three (43.8%) individuals presented with clonality, of whom 26 (60.5%) with minor IGH clonotypes and 17 (39.5%) with major IGH clonotypes. Fifty-five individuals did not present with BCR IGH clonality at any timepoint. Technical replicates were performed for 74 clonal samples from 39 individuals for which sufficient DNA was available to evaluate technical stability of clonality evaluation. All major IGH clonotypes remained clonal (26 out of 26 samples, supplementary figure 1A), while minor IGH clonotypes were more unstable (31 out of 48 remained clonal, 64.6%, supplementary figure 1B). Among the 57 samples with concordant BCR IGH clonality, the clonality category (major vs. minor) was inconsistent between repeats for 5 minor IGH clonotypes and 2 major IGH clonotypes. Diversity among the repeated clonal samples remained relatively stable (mean delta = 0.15, SD = 0.28, supplementary figure 1C).

**Figure 2.**
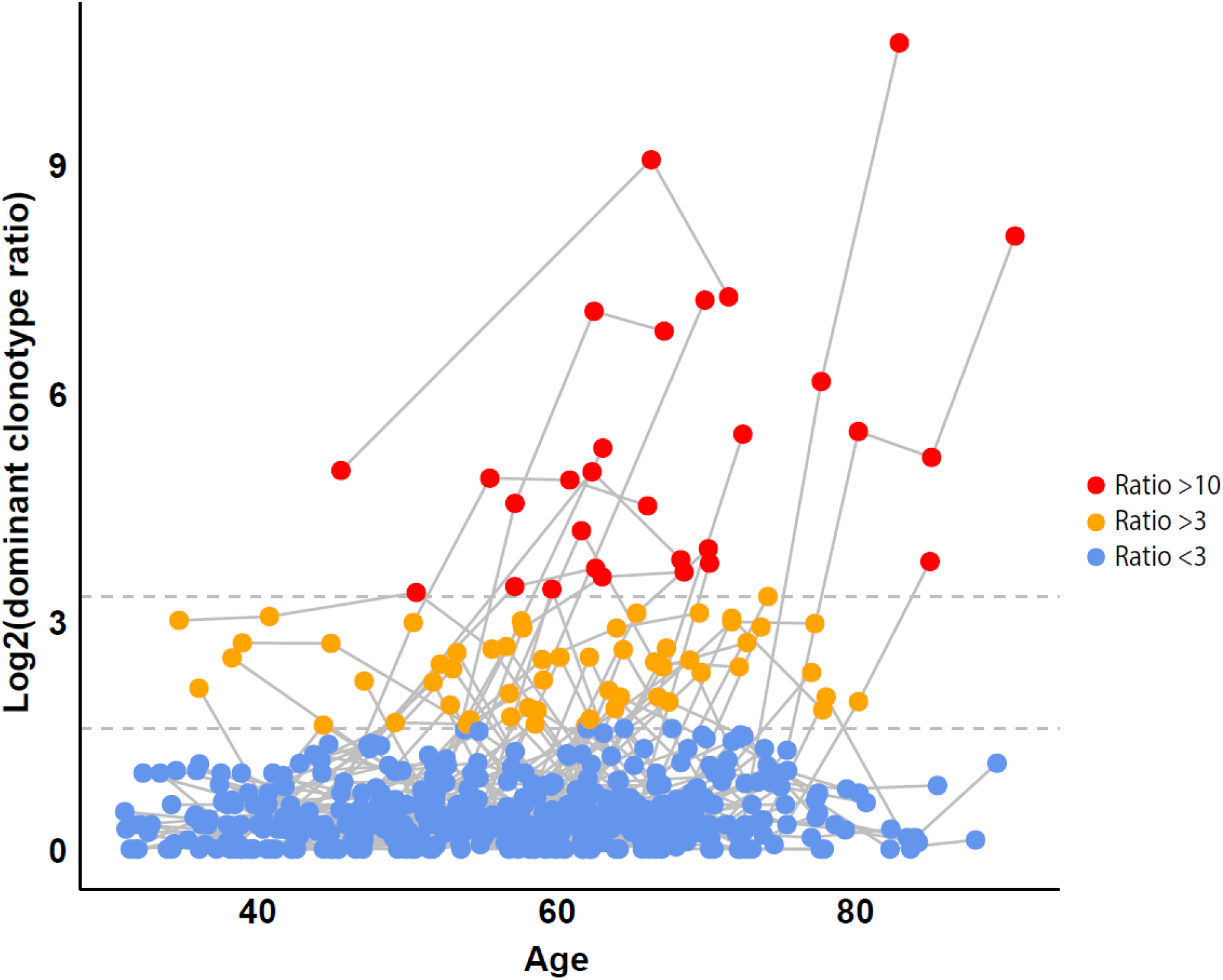
BCR IGH clonality dynamics during aging. Participants with a dominant clonotype ratio between 3 and 10 were designated as “minor IGH clonotypes” and labeled in orange, while individuals with a clonality ratio exceeding 10 were designated as “major IGH clonotypes” and labeled in red. For further details, see methods. Each dot represents a sample, and each observation from the same individual is connected with a line. Number of participants: 98. Total number of observations: 548. X-axis: age at blood draw in years. Y-axis: dominant clonotype ratio with a log2 scale.

Persistence of clonality was evaluated for those individuals for whom at least one follow-up sample was available after the sample in which BCR IGH clonality was first encountered (N = 40). Major IGH clonotypes were significantly more likely to persist and remain clonal than minor IGH clonotypes (90.9% vs. 31%, Supplementary Table 3, p = 0.006). Minor IGH clonotypes were frequently transient and regressed within 5 years in 20 individuals (69%). Major IGH clonotypes persisted for over 20 years in multiple individuals. We employed a linear mixed-effects model adjusted for sex, smoking status, and alcohol consumption to quantify the effect of 10-year age categories on the log^2^-transformed dominant clonotype ratio. The model revealed that the log_2_-transformed dominant clonotype ratio was significantly increased among participants in age categories 60-70 (β =0.59, p < 0.001) and 70+ (β =0.94, p < 0.0001, Supplementary Table 4).

### Somatic hypermutation status of persistently clonal IGH clonotypes

Somatic hypermutation of the BCR is known to significantly alter its affinity for antigens and may, in some cases, contribute to the pathogenesis of B-cell malignancies.^28^ In our current study, we observed 19 individuals presenting with persistent BCR IGH clonality (supplementary table 3). The somatic hypermutation status of these IGH clonotypes was generally stable, with 16 out of 19 participants (84.2%) presenting with the exact same SHM status at all measurements in which they were clonal (Supplementary Table 5).

### Relationship between aging, IGH gene repertoire diversity and BCR IGH clonality

Our beforementioned findings raise the question whether the observed reduction in IGH gene repertoire diversity with age is driven by the observed increased incidence of BCR IGH clonality reminiscent of MBL. To address this question, we employed a linear mixed-effects model with 10-year age categories, clonality category, sex, smoking status, and alcohol consumption as fixed effects and subject ID as a random effect. The model revealed that IGH gene repertoire diversity was decreased for individuals with major clonotypes (dominant clonotype ratio above 10, β =-0.57, p < 0.001), but not for individuals with minor clonotypes (dominant clonotype ratio between 3 and 10, β =0.14, p = 0.18, Table 2). We observed a trend for a slight decrease in IGH gene repertoire diversity in individuals aged 60-70 (β =-0.19, p = 0.1) and participants over age 70 (β =-0.20, p= 0.13). Therefore, the observed reduction in IGH gene repertoire diversity with older age can partially be explained by the increased incidence of major BCR IGH clonality.

**Table 2.** Results from linear mixed effects model 3.

| Parameter | $\beta$ | 95% CI | P-value |
| --- | --- | --- | --- |
| Intercept | 8.07 | [7.75, 8.38] | <0.0001 |
| Ratio >10 | -0.57 | [-0.87, -0.28] | <0.001 |
| Ratio > 3 and < 10 | 0.14 | [-0.06, 0.35] | 0.18 |
| Age category 40-50 | -0.11 | [-0.33, 0.11] | 0.34 |
| Age category 50-60 | -0.12 | [-0.34, 0.09] | 0.26 |
| Age category 60-70 | -0.19 | [-0.41, 0.03] | 0.10 |
| Age category 70+ | -0.20 | [-0.46, 0.06] | 0.13 |
| Sex | 0.14 | [-0.06, 0.35] | 0.18 |
| Former smoker | 0.02 | [-0.21, 0.26] | 0.88 |
| Non-smoker | -0.04 | [-0.31, 0.24] | 0.79 |
| Alcohol consumption (glasses per day) | 0.04 | [-0.04, 0.11] | 0.33 |

Note: A dominant clonotype ratio is calculated by dividing the frequency of the most dominant BCR clonotype by the mean of the frequency of the clonotypes ranked 3<sup>rd</sup>-7<sup>th</sup>.

This model includes IGH gene repertoire diversity as quantified by the Shannon’s diversity index as the dependent variable, clonality category (dominant clonotype ratio >10, dominant clonotype ratio between 3 and 10, and dominant clonotype ratio <3), age category, sex, smoking status and alcohol consumption as fixed effects and participant ID as a random effect.

### Relationship between B-cell count, IGH gene repertoire diversity and BCR IGH clonality

B-cell counts were measured through flow cytometry for round 6 and 7 of the Doetinchem study. B-cell frequency ranged from 0.4% to 10.3% of total leukocytes (median 2.7%). To evaluate the relationship between IGH gene repertoire diversity, dominant clonotype ratio and B-cell count, we next employed a linear mixed effects model with B-cell count per microliter blood as the dependent variable, IGH gene repertoire diversity, clonality category, 10-year age categories, sex and smoking status as fixed effects and subject ID as a random effect. B-cell counts were increased among individuals with an increased IGH gene repertoire diversity (β =34.5, p=<0.001). In alignment with these findings, we observed a strong association between B-cell template input quantified through spike-in during sequencing and IGH gene repertoire diversity (supplementary figure 2). Participants with both major (β =46.9, p=0.04, Supplementary Table 6) and minor clonotypes (β =38.6, p=0.03) presented with increased B-cell counts. Women additionally had higher B-cell counts than men (β =44.9, p=0.01). Importantly however, no evidence of lymphocytosis (>4000 lymphocytes per microliter blood) was observed for any of the participants.

### Relationship between B-cell subsets and IGH gene repertoire diversity and BCR IGH clonality

The available flow cytometry panel allowed for differentiation of B-cell subsets, namely transitional B-cells, naïve B-cells, switched memory B-cells, non-switched memory B-cells and double-negative (IgD-, CD27-) B-cells in sampling rounds 6 and 7 of the study. In round 7 of the study two additional markers were added to the flowcytometry panel allowing for the identification of age-associated (IgD-, CD27-, CD21+, CD11c-) B-cells. Of these B-cell subsets, only transitional B-cells and naïve B-cells were significantly associated with IGH gene repertoire diversity (Figure 3AB, Supplementary tables 7 and 8). In contrast, BCR IGH clonality was associated with increased counts of age-associated B-cells (Figure 3C, supplementary table 9) and switched memory B-cells (supplementary table 10).

**Figure 3.**
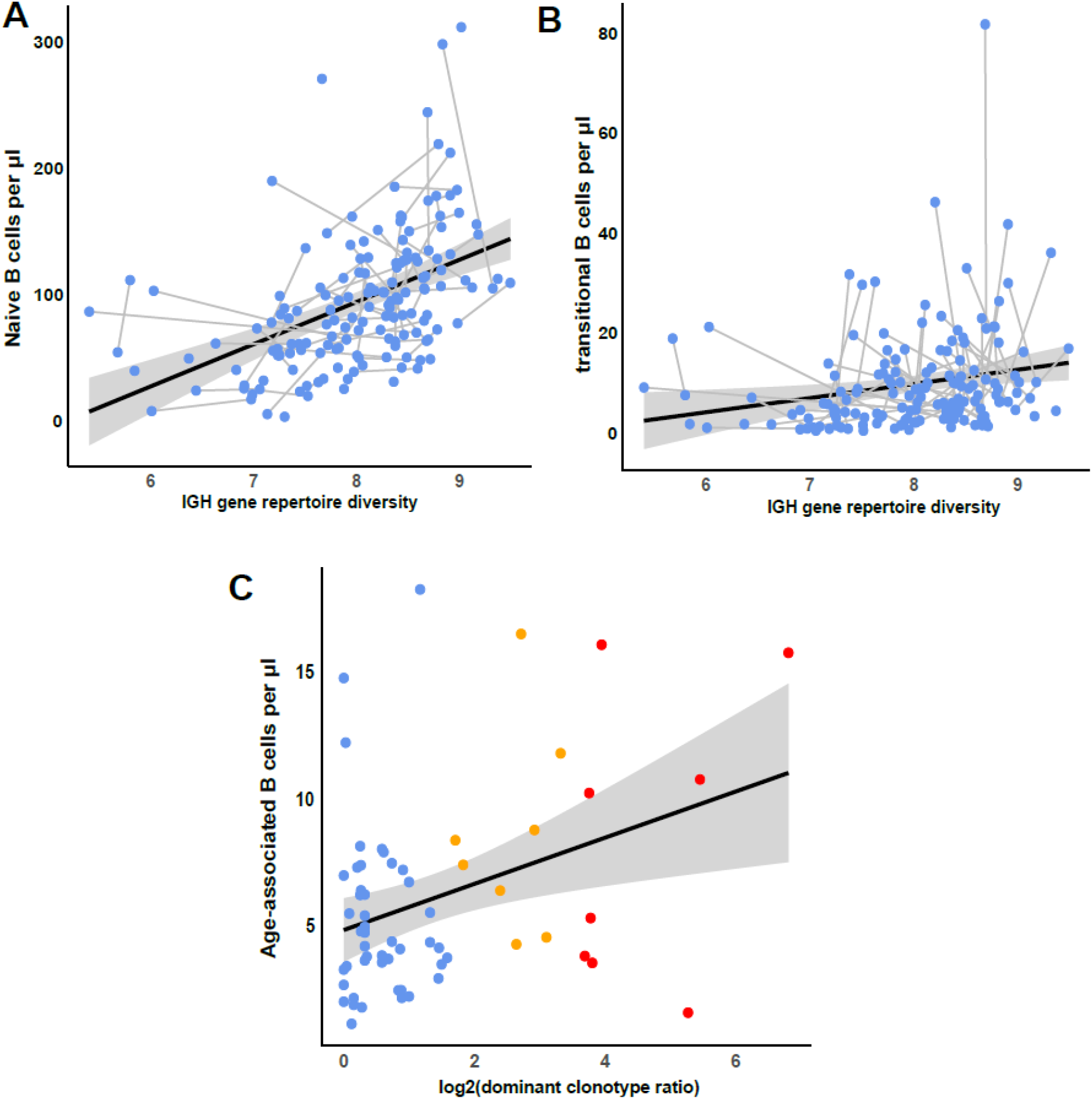
Comparison of B-cell subset counts measured through flow cytometry to IGH gene repertoire diversity and BCR IGH clonality. The black line represents the linear regression estimate that summarizes the general trend. Each dot represents a sample, and each observation from the same individual is connected with a line. A) Naïve B-cel counts per microliter blood versus IGH gene repertoire diversity. B) Transitional B-cell counts per microliter blood versus IGH gene repertoire diversity. C) Age-associated B-cell counts per microliter blood versus log2-transformed dominant clonotype ratio as a proxy for IGH gene repertoire clonality. Participants with a dominant clonotype ratio between 3 and 10 were designated as “minor IGH clonotypes” and labeled in orange, while individuals with a clonality ratio exceeding 10 were designated as “major IGH clonotypes” and labeled in red.

### Relationship between infectious titers and BCR IGH clonality

Infectious titers were measured for VZV, EBV and CMV in round 6 of the Doetinchem study. To evaluate whether increased viral titers may explain the incidence of BCR IGH clonality, we employed a logistic regression model with clonality category as the outcome variable and VZV, EBV and CMV concentration, sex and 10-year age categories as predictors. The model did not reveal any significant associations between clonality category and viral titers (Supplementary Table 11).

## Discussion

In this longitudinal study evaluating dynamics of the IGH gene repertoire during aging over a 30 year period in 98 adults, we observed a notable decrease in IGH gene repertoire diversity starting around age 60. This reduction in diversity coincided with an increased incidence of BCR IGH clonality. Increased IGH gene repertoire diversity was associated with increased transitional and naïve B-cell counts, while BCR IGH clonality was associated with increased counts of non-switched memory B-cells and age-associated B-cells.

The alterations in the IGH gene repertoire observed starting around age 60 are consistent with previous reports describing the emergence of immunosenescence in the 6^th^ decade of life. Indeed, a previous spectratyping-based study in individuals aged 86-94 years old revealed a significant reduction in IGH gene repertoire diversity in this very elderly population compared to younger healthy controls aged 19-54 years.^2,3,29,30^ This reduced IGH gene repertoire diversity may contribute to the impaired humoral immunity associated with aging and may reflect impaired B-cell development in the bone marrow, resulting in the accumulation of atypical B-cells, referred to as age-associated B cells.^31^ Age-associated B-cells have an atypical immunophenotype and are associated with various auto-immune diseases through autoantibody production, inflammatory cytokine production and dysregulated T-cell interactions.^32^ Indeed, we observed that IGH gene repertoire clonality coincided with increased counts of age-associated B-cells.

These age-related changes in the B-cell compartment pose a puzzle to clinicians who want to evaluate the immunological health of their aging patients. In this aging population, incidental findings of B-cell clonality in the peripheral blood during routine checkups are common.^33^ It is frequently unclear whether such findings represent early stages of lymphoid malignancy (*e*.*g*. MBL) or merely benign, transient clonal expansions. Based on the high frequency and temporal and technical instability of the low-level clonality we observed among individuals recruited from the general population, we suggest using a conservative dominant clonotype ratio cutoff of 10 to assess BCR IGH clonality in the peripheral blood and a watch and wait approach for incidental clonal findings. A single follow up measurement performed one to two years after the incidental finding may reveal regression of BCR IGH clonality for some affected individuals and progression for others, meaning longitudinal monitoring is essential in this context.

In the event that the clonality in the peripheral blood turns out to be persistent, an individual may be diagnosed with MBL. MBL is generally split into two categories: low count and high count. Low count MBL is defined as fewer than 500 monoclonal B-cells per microliter blood. Risk of progression to lymphoid malignancy is lower for low-count MBL (4.3-fold increased risk compared to controls), than for high-count MBL (74-fold increased risk).^12^ The lack of lymphocytosis and the weak association between B-cell count and dominant clonotype ratio observed in this study highlight that low-level BCR IGH clonality dynamics may have minimal impact on total B-cell count.^34^ Like MBL, BCR IGH clonality is associated with lymphoid malignancy risk.^34^

Historically, common infections like pneumonia and herpes zoster have been suggested as potential causal triggers of MBL and CLL development.^35-37^ Consistent with this hypothesis, vaccination has been suggested to reduce MBL and CLL risk.^38,39^ Notably, individuals with MBL are also at increased risk of developing subsequent infections.^40,41^ However, a recent study in a cohort of 1009 MBLs and 4419 controls showed no evidence for infectious disease as a risk factor for MBL and no evidence for vaccination as a protective risk factor.^42^ These recent findings are consistent with our current study, where VZV, EBV and CMV titers did not contribute to BCR IGH clonality risk. However, viral titers were only available for one out 7 sampling rounds in our current study. Further research is therefore required to evaluate whether BCR IGH clonality may predispose individuals to infections, as has been shown for low-count MBL.

MBL is traditionally diagnosed using flow cytometry. A notable limitation of our study is that flow cytometry data was only available for round 6 and 7 of the study and viral titers only in round 6. Additionally, the antibody panel used for flow cytometry was not optimized for MBL diagnosis, but rather focused on providing an overview of the major cell lineages. Therefore, a longitudinal validation study combining flowcytometry-based and molecular-based MBL diagnostics is a crucial next step to validate our molecular approach to MBL diagnosis. Another limitation included the DNA input of 500 ng, translating to a median input of 3447 B-cells (range 57-17,361). This relatively shallow pool of sequenced B-cells naturally results in an underestimation of the B-cell diversity, which may limit the generalizability of our findings. UMI-based error correction would additionally be helpful to explore the degree to which PCR errors may be affecting diversity and clonality estimates.

In our study, BCR IGH clonality was unstable for around one third of minor IGH clonotypes in technical replicates. Although this instability was caused by relatively minor shifts in abundance ratio close to the clonality cutoff for some samples, others showed a more pronounced change in abundance ratio, highlighting that technical variance in the form of sampling bias or amplification bias can result in inaccurate clonality assessment, particularly for minor clonotypes. This artificial clonality has previously been described in the literature and is sometimes referred to as pseudoclonality. The detected instability may be partially mitigated by increasing the DNA input of the assay to expand the B-cell pool that is being measured. The risk of pseudoclonality further exemplifies the need to use a conservative dominant clonotype ratio cutoff of 10.

Previous longitudinal studies were limited in the number of individuals included and the length of follow up.^43^ Their focus also tended to be on the composition and differentiation of immune cell subsets, such as memory B-cells and plasmablasts, rather than on the overall clonality and diversity of the IGH gene repertoire. Our current study extends these findings through the long follow up duration (30 years) and high sampling frequency (every 5 years).

## Conclusions

In summary, our study indicates that IGH gene repertoire diversity decreases significantly after age 60 and that BCR IGH clonality becomes increasingly common during ageing. IGH gene repertoire diversity was determined primarily by the naïve and transitional B-cell pool, while BCR IGH clonality was influenced by switched memory and age-associated B-cell counts. Major IGH clonotypes with a dominant clonotype ratio above 10 are significantly more likely to persist than minor IGH clonotypes below this cutoff. Longitudinal analyses such as the present study provide valuable insight into the dynamic nature of IGH gene repertoire remodeling during aging. Future research integrating multi-omics and single-cell approaches will be essential to further elucidate the biological mechanisms underlying age-related clonal expansions and their role in the development of lymphoid malignancies.

## Acknowledgements

The Doetinchem Cohort Study is supported by the Dutch Ministry of Health, Welfare and Sport and the National Institute for Public Health and the Environment. We thank the respondents, epidemiologists, and fieldworkers of the Municipal Health Service in Doetinchem for their contribution to the data collection for this study. In particular, the authors thank Annemarie Teitsema-Jansen and Petra Vissink for coordinating sample retrieval for the project.

## Contribution

M.V. contributed to the development and maintenance of the Doetinchem Cohort Study. A.B. and M.Z.B. developed and implemented the DCS subcohort study design. L.L. and M.A.J.I.O. performed the immunogenetic sequencing experiments. M.Z.B. and L.G.H.R. developed the flow cytometry laboratory protocols and performed the flow cytometry experiments. L.L. and P.M.K. analyzed the data. L.L. and P.M.K. wrote the manuscript. M.A.J.I.O., P.E., A.B., M.Z.B., L.R., W.M., R.C.H.V., and A.W.L. critically reviewed and edited the manuscript.

## Ethics approval and consent to participate

The study was approved by the local institutional medical ethical committee at the Erasmus MC under protocol MEC 2019-0484. All participants written informed consent for every DCS round and for this subcohort study separately. The study was performed in compliance with the declaration of Helsinki.

### Abbreviations

BCR IGH: B-cell receptor immunoglobulin heavy chain
IGHV: immunoglobulin heavy chain variable
VZV: Varicella Zoster virus
CMV: Cytomegalovirus
EBV: Epstein-Barr virus
SASP: senescence-associated phenotype
MBL: monoclonal B-cell lymphocytosis
CLL: chronic lymphocytic leukemia
CDR3: complementarity-determining region 3
LOESS: Locally Estimated Scatterplot Smoothing

## Consent for publication

Not applicable

## Availability of data and materials

The datasets used are available from the corresponding author upon reasonable request and with permission of the scientific committee of the Doetinchem Cohort Study.

