## Supplementary material for "Longitudinal analysis of the B-cell receptor repertoire across 30 years of ageing"

**Supplementary Table 1** - A list of the antibodies and additional reagents used within this study to immunophenotype the repeated blood samples of the study participants collected in 2024.

| Antigen | Fluorochrome | Clone | Company | Catalogue # |
|---|---|---|---|---|
| **CD45** | RealBlue613 | HI30 | BD Biosciences | 758752 |
| **CD3** | BV605 | UCHT1 | Biolegend | 300460 |
| **CD19** | BUV395 | HIB19 | BD Biosciences | 569107 |
| **CD8a** | FITC | RPA-T8 | Biolegend | 301006 |
| **CD14** | PerCP-Cy5.5 | MoP9 | BD Biosciences | 562692 |
| **CD127** | APC | A019D5 | Biolegend | 351316 |
| **CXCR5** | APC-R700 | RF8B2 | BD Biosciences | 565191 |
| **IgD** | BV605 | IA6-2 | Biolegend | 348232 |
| **HLA-DR** | BV650 | G46-6 | BD Biosciences | 564231 |
| **CD27** | BV711 | O323 | Biolegend | 302834 |
| **CD16** | BV786 | 3G8 | BD Biosciences | 563690 |
| **CD38** | PE-CF594 | HIT2 | BD Biosciences | 562288 |
| **CD25** | PE-Cy7 | M-A251 | BD Biosciences | 557741 |
| **CD56** | BUV563 | NCAM16.2 | BD Biosciences | 612928 |
| **CD4** | BUV615 | SK3 | BD Biosciences | 612987 |
| **CD45RA** | BUV737 | HI100 | BD Biosciences | 612846 |
| **CD28** | **RB744** | CD28.2 | BD Biosciences | 758130 |
| **CD21** | **BV480** | B-ly4 | BD Biosciences | 746613 |
| **CD11c** | **BV750** | B-ly6 | BD Biosciences | 747459 |
| **True-Stain Monocyte blocker** | | | Biolegend | 426102 |
| **TruStain FcX** | | | Biolegend | 422302 |

**Supplementary Table 2** – Results from linear mixed effects model 1. This model includes IGH gene repertoire diversity as quantified by the Shannon's diversity index as the dependent variable, age category, sex, smoking status and alcohol consumption as fixed effects and participant ID as a random effect.

| Parameter | β | 95% CI | P-value |
|---|---|---|---|
| Intercept | 8.10 | [7.78, 8.42] | <0.0001 |
| Age category 40-50 | -0.13 | [-0.35, 0.10] | 0.28 |
| Age category 50-60 | -0.14 | [-0.36, 0.07] | 0.19 |
| **Age category 60-70** | -0.24 | [-0.46, -0.02] | **0.03** |
| **Age category 70+** | -0.27 | [-0.53, -0.01] | **0.04** |
| Sex | 0.13 | [-0.09, 0.35] | 0.24 |
| Former smoker | 0.02 | [-0.22, 0.27] | 0.88 |

| | | | |
|---|---|---|---|
| Non-smoker | -0.03 | [-0.31, 0.26] | 0.86 |
| Alcohol consumption (glasses per day) | 0.03 | [-0.04, 0.10] | 0.43 |

**Supplementary Table 3** – Clonality persistence. Clonotypes with a dominant clonotype ratio between 3 and 10 were designated as "minor clonotypes", while clonotypes with a dominant clonotype ratio exceeding 10 were designated as "major clonotypes". A clonotype was designated as "persistent" if it reached the cutoff for clonality in two sequential samples drawn 5 years apart. In total, 40 individuals presented with at least one minor or major clonotype with a follow up sample available to evaluate persistence.

| | Minor (n = 29) | Major (n = 11) | Total (n = 40) |
|---|---|---|---|
| Persistent N (%) | 9 (31%) | 10 (90.9%) | 19 (47.5%) |
| Regressing N (%) | 20 (69%) | 1(9.1%) | 21 (52.5%) |

**Supplementary Table 4** – Results from linear mixed effects model 2. This model includes the $log_2$-transformed dominant clonotype ratio as the dependent variable, age category, sex, smoking status and alcohol consumption as fixed effects and participant ID as a random effect.

| Parameter | β | 95% CI | P-value |
|---|---|---|---|
| Intercept | 0.51 | [-0.02, 0.99] | 0.06 |
| Age category 40-50 | 0.07 | [-0.29, 0.36] | 0.72 |
| Age category 50-60 | 0.23 | [-0.17, 0.44] | 0.18 |
| **Age category 60-70** | 0.59 | [0.16, 0.79] | **<0.001** |
| **Age category 70+** | 0.94 | [0.48, 1.21] | **<0.0001** |
| Sex | -0.04 | [-0.41, 0.30] | 0.85 |
| Former smoker | 0.09 | [-0.37, 0.33] | 0.67 |
| Non-smoker | -0.10 | [-0.62, 0.25] | 0.68 |
| Alcohol consumption (glasses per day) | 0.04 | [-0.08, 0.14] | 0.57 |

**Supplementary Table 5** – Samples with stable clonality across rounds. Germline identity of the IGHV gene is shown. While the top clonotype for participant 4 shifted in round 7 to a newly emerged clonotype, the previous clonotype was still ranked second and remained clonal with stable SHM status (data not shown).

| ID | Round | Ratio | diversity | log2_ratio | Age | Clonality | Germline identity | Vgene | CDR3 |
|---|---|---|---|---|---|---|---|---|---|
| 1 | DCS5 | 5.4 | 8.22 | 2.43 | 52.19 | Ratio >3 | 95.83% | V1-69 | CAREGRSGVTNPIDYW |
| 1 | DCS6 | 10.95 | 8.67 | 3.45 | 57.19 | Ratio >10 | 95.83% | V1-69 | CAREGRSGVTNPIDYW |
| 1 | DCS7 | 12.93 | 8.3 | 3.69 | 62.60 | Ratio >10 | 95.83% | V1-69 | CAREGRSGVTNPIDYW |
| 2 | DCS5 | 5.63 | 7.93 | 2.5 | 59.03 | Ratio >3 | 96.53% | V3-23 | CAKEDLQHSWARCW |
| 2 | DCS6 | 7.5 | 7.96 | 2.91 | 64.01 | Ratio >3 | 92.01% | V3-21 | CAREADDGDSRRPFDPW |
| 2 | DCS7 | 8.6 | 7.83 | 3.1 | 69.56 | Ratio >3 | 92.01% | V3-21 | CAREADDGDSRRPFDPW |
| 3 | DCS6 | 8 | 8.65 | 3 | 57.62 | Ratio >3 | 91.67% | V3-30 | CAKVINGGEFRDYRYYGMDAW |
| 3 | DCS7 | 38.7 | 7.72 | 5.27 | 63.09 | Ratio >10 | 91.67% | V3-30 | CAKVINGGEFRDYRYYGMDAW |
| 4 | DCS5 | 45 | 8.13 | 5.49423 | 80.22 | Ratio >10 | 98.28% | V4-61 | CARAGGVGYRYASQPPRRGNYNWFDPW |

| | | | | | | | | | |
|---|---|---|---|---|---|---|---|---|---|
| 4 | DCS6 | 35.50 | 7.66 | 5.15 | 85.12 | Ratio >10 | 98.28% | V4-61 | CARAGGVGYRYASQPPRRGNYNWFDPW |
| 4 | DCS7 | 267.50 | 4.70 | 8.06 | 90.72 | Ratio >10 | 94.04% | V4-59 | CARDRRNALATHYFDYW |
| 5 | DCS5 | 3.53 | 8.46 | 1.82 | 58.69 | Ratio >3 | 94.44% | V3-7 | CAVAKLQGEYYDYW |
| 5 | DCS7 | 5.00 | 8.33 | 2.32 | 69.68 | Ratio >3 | 94.10% | V3-7 | CAVAKLQGEYYDYW |
| 6 | DCS1 | 8.04 | 8.07 | 3.00 | 34.69 | Ratio >3 | 95.14% | V1-69 | CAREFSYESSGYYYLYW |
| 6 | DCS2 | 8.33 | 7.60 | 3.06 | 40.75 | Ratio >3 | 95.14% | V1-69 | CAREFSYESSGYYYLYW |
| 6 | DCS4 | 10.40 | 7.60 | 3.37 | 50.59 | Ratio >10 | 95.14% | V1-69 | CAREFSYESSGYYYLYW |
| 6 | DCS5 | 6.19 | 7.75 | 2.63 | 55.67 | Ratio >3 | 95.14% | V1-69 | CAREFSYESSGYYYLYW |
| 7 | DCS1 | 31.54 | 7.96 | 4.98 | 45.54 | Ratio >10 | 93.75% | V1-69 | CAREGRQLGTSNSFDYW |
| 7 | DCS5 | 535.80 | 5.86 | 9.07 | 66.34 | Ratio >10 | 93.75% | V1-69 | CAREGRQLGTSNSFDYW |
| 7 | DCS6 | 153.50 | 6.37 | 7.26 | 71.52 | Ratio >10 | 93.75% | V1-69 | CAREGRQLGTSNSFDYW |
| 8 | DCS5 | 7.50 | 7.54 | 2.91 | 57.73 | Ratio >3 | 98.95% | V4-34 | CARVQDSSGYFAFDIW |
| 8 | DCS6 | 11.96 | 7.52 | 3.58 | 63.05 | Ratio >10 | 98.95% | V4-34 | CARVQDSSGYFAFDIW |
| 8 | DCS7 | 12.50 | 8.08 | 3.64 | 68.54 | Ratio >10 | 99.65% | V4-34 | CARVQDSSGYFAFDIW |
| 9 | DCS4 | 7.88 | 8.03 | 2.98 | 50.39 | Ratio >3 | 95.24% | V3-15 | CTTGVQRFLQAGYW |
| 9 | DCS5 | 29.40 | 8.09 | 4.88 | 55.52 | Ratio >10 | 95.24% | V3-15 | CTTGVQRFLQAGYW |
| 9 | DCS6 | 28.92 | 7.51 | 4.85 | 60.88 | Ratio >10 | 95.24% | V3-15 | CTTGVQRFLQAGYW |
| 9 | DCS7 | 22.86 | 8.04 | 4.51 | 66.09 | Ratio >10 | 95.24% | V3-15 | CTTGVQRFLQAGYW |
| 10 | DCS6 | 71 | 5.85 | 6.15 | 77.72 | Ratio >10 | 95.58% | V3-73 | CARHTTRGLYW |
| 10 | DCS7 | 1555 | 1.16 | 10.60 | 82.96 | Ratio >10 | 95.58% | V3-73 | CARHTTRGLYW |
| 11 | DCS5 | 7.94 | 9.10 | 2.99 | 71.73 | Ratio >3 | 96.84% | V4-34 | CARYFPDCSTTSCYEPTRHYYYYYMDVW |
| 11 | DCS6 | 7.81 | 8.81 | 2.97 | 77.30 | Ratio >3 | 96.84% | V4-34 | CARYFPDCSTTSCYEPTRHYYYYYMDVW |
| 12 | DCS5 | 23.38 | 8.72 | 4.55 | 57.20 | Ratio >10 | 98.96% | V3-48 | CARAGNYDDHYYFMDVW |
| 12 | DCS6 | 134.67 | 5.80 | 7.07 | 62.50 | Ratio >10 | 98.96% | V3-48 | CARAGNYDDHYYFMDVW |
| 12 | DCS7 | 112.34 | 5.80 | 6.81 | 67.20 | Ratio >10 | 98.96% | V3-48 | CARAGNYDDHYYFMDVW |
| 13 | DCS6 | 3.83 | 7.70 | 1.94 | 80.22 | Ratio >3 | 92.71% | V3-11 | CARVSGYSYGYADRDYDYYYYMDVW |
| 13 | DCS7 | 13.75 | 7.82 | 3.78 | 85.02 | Ratio >10 | 92.71% | V3-11 | CARVSGYSYGYADRDYDYYYYMDVW |
| 14 | DCS6 | 8.59 | 8.02 | 3.1 | 65.36 | Ratio >3 | 98.25% | V3-66 | CARTTSGRRASRGDCYAPRNGMDVW |
| 14 | DCS7 | 15.45 | 8.08 | 3.95 | 70.15 | Ratio >10 | 98.25% | V3-66 | CARTTSGRRASRGDCYAPRNGMDVW |
| 15 | DCS6 | 5.26 | 9.12 | 2.39 | 67.1 | Ratio >3 | 97.54% | V4-34 | CARPLGRRGSWFDPW |
| 15 | DCS7 | 6.57 | 8.27 | 2.72 | 72.74 | Ratio >3 | 97.54% | V4-34 | CARPLGRRGSWFDPW |
| 16 | DCS4 | 4.13 | 8.69 | 2.05 | 56.83 | Ratio >3 | 93.33% | V4-34 | CASAGAAGMRWFDPW |
| 16 | DCS5 | 18.22 | 8.59 | 4.18 | 61.65 | Ratio >10 | 96.14% | V4-34 | CASAGAAGMRWFDPW |
| 16 | DCS6 | 5.5 | 7.17 | 2.46 | 66.56 | Ratio >3 | 91.67% | V3-30 | CAKVINGGEFRDYRYYGMDAW |
| 16 | DCS7 | 5.26 | 7.5 | 2.40 | 72.22 | Ratio >3 | 94.04% | V4-34 | CASAGAAGMRWFDPW |
| 17 | DCS4 | 3.13 | 8.29 | 1.65 | 54.00 | Ratio >3 | 99.31% | V3-21 | CARGYNYGYFTRYYYYGMDVW |
| 17 | DCS5 | 4.67 | 8.05 | 2.22 | 59.1 | Ratio >3 | 99.65% | V3-21 | CARGYNYGYFTRYYYYGMDVW |
| 17 | DCS7 | 149.14 | 7.83 | 7.22 | 69.30 | Ratio >10 | 99.65% | V3-21 | CARGYNYGYFTRYYYYGMDVW |
| 18 | DCS6 | 31.18 | 7.45 | 4.96 | 62.37 | Ratio >10 | 98.26% | V3-7 | CARGLPHNIVVGEVQGSW |
| 18 | DCS7 | 14 | 7.63 | 3.81 | 68.32 | Ratio >10 | 98.26% | V3-7 | CARGLPHNIVVGEVQGSW |
| 19 | DCS6 | 4 | 8.48 | 2 | 66.78 | Ratio >3 | 89.24% | V3-23 | CAKEGGSTISSLGQW |
| 19 | DCS7 | 43.93 | 8.47 | 5.46 | 72.48 | Ratio >10 | 89.24% | V3-23 | CAKEGGSTISSLGQW |

**Supplementary Table 6** – Results from linear mixed effects model 4. This model includes B-cell count per microliter blood as the dependent variable, IGH gene repertoire diversity, clonality category (dominant clonotype ratio >10, between 3 and 10 or <3), age in years, sex and smoking status and alcohol consumption as fixed effects and participant ID as a random effect. B-cell count data were only available for a total of 156 samples from round 6 and 7 of the study.

| **Parameter** | **β** | **95% CI** | **P-value** |
|---|---|---|---|
| Intercept | -232 | [-456, -14.9] | **0.04** |
| Ratio >10 | 46.9 | [4.3, 90.9] | **0.04** |
| Ratio >3 and < 10 | 38.6 | [4.9, 72.5] | **0.03** |
| IGH gene repertoire diversity | 34.5 | [16.3, 53.9] | **<0.001** |
| Age | 1.2 | [-0.8, 3.2] | 0.25 |
| Sex (female) | 44.9 | [10.5, 72.1] | **0.01** |
| Former smoker | 22.1 | [-40.3, 84.5] | 0.5 |
| Non-smoker | 21.5 | [-43.9, 87.0] | 0.53 |
| Alcohol consumption (glasses per day) | -0.61 | [-13.4, 12.2] | 0.93 |

**Supplementary Table 7** – Results from linear mixed effects model 5. This model includes Naïve B-cell count per microliter blood as the dependent variable, IGH gene repertoire diversity, clonality category (dominant clonotype ratio >10, between 3 and 10 or <3), age in years, sex and smoking status and alcohol consumption as fixed effects and participant ID as a random effect. Naïve B-cell count data were only available for a total of 156 samples from round 6 and 7 of the study.

| **Parameter** | **β** | **95% CI** | **P-value** |
|---|---|---|---|
| Intercept | -210 | [-355, -71.6] | **<0.01** |
| Ratio >10 | 5.3 | [-21.2, 32.5] | 0.71 |
| Ratio >3 and < 10 | 1.54 | [20.4, 23.2] | 0.89 |
| IGH gene repertoire diversity | 28.35 | [17.2, 40.2] | **<0.0001** |
| Age | 0.61 | [-0.6, 1.9] | 0.34 |
| Sex (female) | 8.6 | [-10.3, 27.4] | 0.39 |
| Former smoker | 33.2 | [-5.4, 72.4] | 0.1 |
| Non-smoker | 33.0 | [-7.1, 73.5] | 0.12 |
| Alcohol consumption (glasses per day) | -0.96 | [-8.8, 6.8] | 0.81 |

**Supplementary Table 8** – Results from linear mixed effects model 6. This model includes transitional B-cell count per microliter blood as the dependent variable, IGH gene repertoire diversity, clonality category (dominant clonotype ratio >10, between 3 and 10 or <3), age in years, sex and smoking status and alcohol consumption as fixed effects and participant ID as a random effect. Transitional B-cell count data were only available for a total of 156 samples from round 6 and 7 of the study.

| Parameter | β | 95% CI | P-value |
|---|---|---|---|
| Intercept | -51.2 | [-77.7 , 23.7 ] | **<0.001** |
| Ratio >10 | 3.2 | [-2.1, 8.6] | 0.25 |
| Ratio >3 and < 10 | -0.54 | [-5, 4.1] | 0.84 |
| IGH gene repertoire diversity | 3.5 | [1.4, 5.7] | **<0.01** |
| Age | 0.39 | [0.12, 0.66] | **<0.01** |
| Sex (female) | 0.87 | [-4.3, 2.6] | 0.63 |
| Former smoker | 7.3 | [-0.7, 15.4] | 0.09 |
| Non-smoker | 5.8 | [-2.4, 14] | 0.18 |
| Alcohol consumption (glasses per day) | -0.96 | [-2.3, 0.73] | 0.33 |

**Supplementary Table 9** – Results from linear regression model 1. This model includes age-associated B-cell count per microliter blood as the dependent variable, log2-transformed dominant clonotype ratio, age in years, sex, smoking status and alcohol consumption as independent variables. Age-associated B-cell count data were only available for a total of 62 samples from round 7 of the study.

| Parameter | β | 95% CI | P-value |
|---|---|---|---|
| Intercept | 8.8 | [-3.5 , 21.1 ] | 0.15 |
| Log2 dominant clonotype ratio | 0.95 | [0.3, 1.6] | **<0.01** |
| IGH gene repertoire diversity | -0.9 | [-2.1,0.3] | 0.13 |
| Age | -0.06 | [-0.23, 0.12] | 0.51 |
| Sex (female) | 1.17 | [-0.95, 3.3] | 0.27 |
| Non-smoker | -0.93 | [-3.1, 1.2] | 0.38 |
| Alcohol consumption (glasses per day) | -0.15 | [-1.4, 1.1] | 0.81 |

**Supplementary Table 10** – Results from linear mixed effects model 7. This model includes switched memory B-cell count per microliter blood as the dependent variable, IGH gene repertoire diversity, clonality category (dominant clonotype ratio >10, between 3 and 10 or <3), age in years, sex and smoking status and alcohol consumption as fixed effects and participant ID as a random effect. Switched memory B-cell count data were only available for a total of 156 samples from round 6 and 7 of the study.

| Parameter | β | 95% CI | P-value |
|---|---|---|---|
| Intercept | 25.3 | [-77.7 , 23.7 ] | 0.41 |
| Ratio >10 | 14.5 | [-2.1, 8.6] | **0.02** |
| Ratio >3 and < 10 | 15.1 | [-5, 4.1] | **<0.01** |
| IGH gene repertoire diversity | 2.9 | [1.4, 5.7] | 0.24 |
| Age | -0.33 | [0.12, 0.66] | 0.25 |
| Sex (female) | 15 | [-4.3, 2.6] | **<0.001** |
| Former smoker | -5 | [-0.7, 15.4] | 0.58 |
| Non-smoker | -10.4 | [-2.4, 14] | 0.26 |
| Alcohol consumption (glasses per day) | 0.2 | [-2.3, 0.73] | 0.92 |

**Supplementary Table 11** – Results from logistic regression model 6. This model includes clonality category (above or below clonality ratio >10) as the outcome variable, VZV titer, EBV titer, CMV titer, age in years and sex as predictors. Viral titer data were only available for round 6 of the study.

| Parameter | OR | 95% CI | P-value |
|---|---|---|---|
| Intercept | 0.02 | [0, 32] | 0.28 |
| Age | 1.05 | [0.94, 1.18] | 0.36 |
| Sex (female) | 0.7 | [0.89, 3.9] | 0.66 |
| CMV titer | 1 | [0.99, 1] | 0.3 |
| EBV titer | 0.99 | [0.98, 1] | 0.16 |
| VZV titer | 0.91 | [0.55, 1.1] | 0.64 |

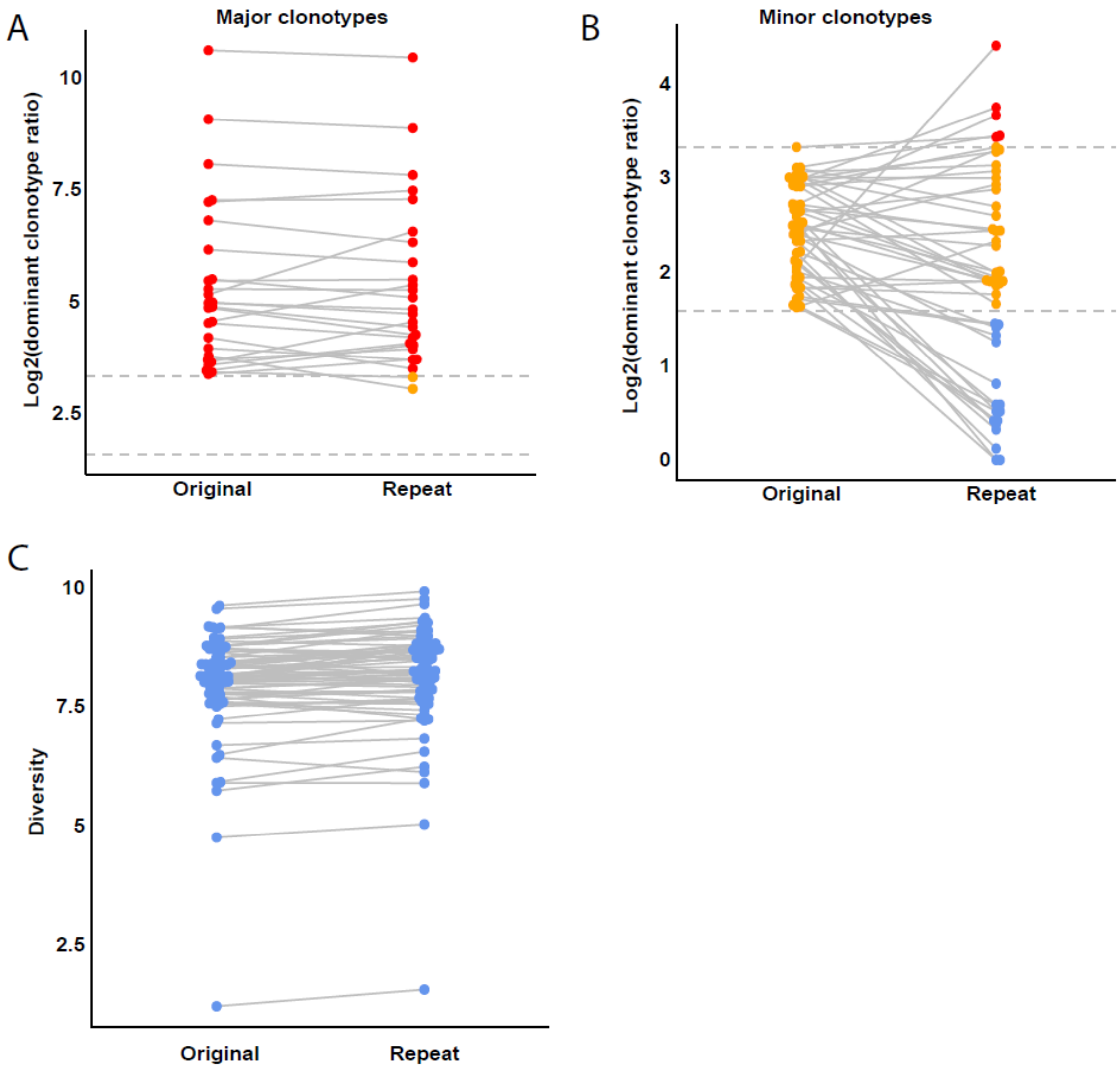


**Supplementary Figure 1. Technical replicates of 74 samples from 39 participants with clonality.** Participants with a dominant clonotype ratio between 3 and 10 were designated as "minor clonotypes" and labeled in orange, while individuals with a clonality ratio exceeding 10 were designated as "major clonotypes" and labeled in red. Paired samples were connected with a line to illustrate dynamics across replicates. A) Stability of the log2-transformed dominant clonotype ratio across replicates for samples with major clonotypes. B) Stability of the log2-transformed dominant clonotype ratio across replicates for samples with minor clonotypes. C) Stability of Shannon diversity index across all technical replicates.

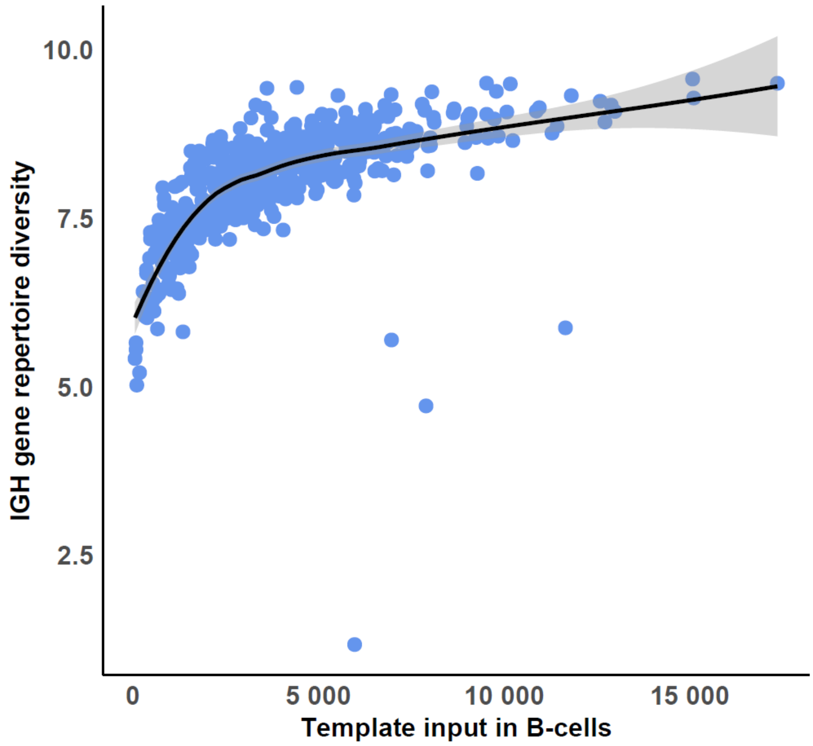


**Supplementary Figure 2. IGH gene repertoire diversity vs. template input in B-cells.** Template input in B-cells was calculated based on spike-in DNA from B-cell cell lines. The black line represents the Locally Estimated Scatterplot Smoothing (LOESS) regression estimate that summarizes the general trend. Outliers represent individuals with increased abundance ratio, meaning individuals with increased IGH gene repertoire clonality.
